# lme4qtl: linear mixed models with flexible covariance structure for genetic studies of related individuals

**DOI:** 10.1101/139816

**Authors:** Andrey Ziyatdinov, Miquel Vázquez-Santiago, Helena Brunel, Angel Martinez-Perez, Hugues Aschard, Jose Manuel Soria

## Abstract

**Background:** Quantitative trait locus (QTL) mapping in genetic data often involves analysis of correlated observations, which need to be accounted for to avoid false association signals. This is commonly performed by modeling such correlations as random effects in linear mixed models (LMMs). The R package *lme4* is a well-established tool that implements major LMM features using sparse matrix methods; however, it is not fully adapted for QTL mapping association and linkage studies. In particular, two LMM features are lacking in the base version of *lme4*: the definition of random effects by custom covariance matrices; and parameter constraints, which are essential in advanced QTL models. Apart from applications in linkage studies of related individuals, such functionalities are of high interest for association studies in situations where multiple covariance matrices need to be modeled, a scenario not covered by many genome-wide association study (GWAS) software.

**Results:** To address the aforementioned limitations, we developed a new R package *lme4qtl* as an extension of *lme4*. First, *lme4qtl* contributes new models for genetic studies within a single tool integrated with *lme4* and its companion packages. Second, *lme4qtl* offers a flexible framework for scenarios with multiple levels of relatedness and becomes efficient when covariance matrices are sparse. We showed the value of our package using real family-based data in the Genetic Analysis of Idiopathic Thrombophilia 2 (GAIT2) project.

**Conclusions:** Our software *lme4qtl* enables QTL mapping models with a versatile structure of random effects and efficient computation for sparse covariances. *lme4qtl* is available at https://github.com/variani/lme4qtl.

## 1 Background

Many genetic study designs induce correlations among observations, including, for example, family or cryptic relatedness, shared environments and repeated measurements. The standard statistical approach used in quantitative trait locus (QTL) mapping is linear mixed models (LMMs), which is able to effectively assess the contribution of an individual genetic locus in the presence of correlated observations [1, 2, 3, 4]. However, LMMs are known to be computationally demanding when applied in large-scale data. Indeed, the LMM approach has the cubic complexity on the sample size per test [3]. This is a major barrier in today’s genome-wide association studies (GWAS), which consist in performing millions of tests in sample size of 10,000 or more individuals. Therefore, recent methodological developments have been focused on reduction in computational cost [4].

There has been a notable improvement for LMMs with a single genetic random effect. Both population based [3, 5, 6] and family-based methods [7] use an initial operation on eigendecomposition of the genetic covariance matrix to rotate the data, thereby removing its correlation structure. The computation time drops down to the quadratic complexity on the sample size per test. When LMMs have multiple random effects, the eigendecomposition trick is not applicable and computational speed up can be achieved by tuning the optimization algorithms, for instance, using sparse matrix methods [8] or incorporating Monte Carlo simulations [9].

However, the decrease in computation time comes at the expense of flexibility. In particular, most efficient LMM methods developed for GWAS assume a single random genetic effect in model specification and support simple study designs, for example, prohibiting the analysis of longitudinal panels (Supplementary Table 1). We have developed a new *lme4qtl* R package that unlocks the well-established *lme4* framework for QTL mapping analysis. Of several existing lme4-based extensions [10, 11, 12], the closest to *lme4qtl* is the *pedigreemm* R package [11]. Although this package does support analysis of related individuals, its main limitation is usage of pedigree annotations rather than custom covariance matrices. Other R packages implement routines to fit LMMs as stand-alone programs, for example, the most recent *sommer* package and references therein [13]. Although a given existing tool might have certain advantages over our package, *lme4qtl* uniquely combines two features, custom covariances and parameter constraints, both of which are integrated in the *lme4* framework (Supplementary Tables 1 and 2).

We demonstrate the computational efficiency and versatility of our package through the analysis of real family-based data from the Genetic Analysis of Idiopathic Thrombophilia 2 (GAIT2) project [14]. More specifically, we first performed a standard GWAS, then showed an advanced model of gene-environment interaction [15], and finally estimated the influence of data sparsity on the computation time.

## 2 Implementation

### 2.1 Linear mixed models

Consider the following polygenic linear model that describes an outcome *y*:

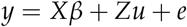

where *n* is the number of individuals, y_n×1_ is vector of size *n*, *X_n×p_* and *Z*_n×n_ are incidence matrices, *p* is the number of fixed effects, *βp×1* is a vector of fixed effects, *u_n×1_* is a vector of a random polygenic effect, and *e_n×1_* is a vector of the residuals errors. The random vectors *u* and *e* are assumed to be mutually uncorrelated and multivariate normally distributed, 𝒩(0, *G_n×n_*) and 𝒩(0, *R_n×n_*). The covariance matrices are parametrized with a few scalar parameters such as 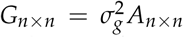. and 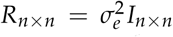, where *A* is a genetic additive relationship matrix and *I* is the identity matrix. In a general case, the model can be extended by adding more random effects, for instance, the dominance genetic or shared-environment components.

### 2.2 Implementation of ***lme4qtl***

As an extension of the *lme4* R package, *lme4qtl* adopts its features related to model specification, data representation and computation [16]. Briefly, models are specified by a single formula, where grouping factors defining random effects can be nested, partially or fully crossed. Also, underlying computation relies on sparse matrix methods and formulation of a penalized least squares problem, for which many optimizers with box constraints are available. While *lme4* fits linear and generalized linear mixed models by means of lmer and glmer functions, *lme4qtl* extends them in `relmatLmer` and `relmatGlmer` functions. The new interface has two main additional arguments: `relmat` for covariance matrices of random effects and vcControl for restrictions on variance component model parameters. Since the developed `relmatLmer` and `relmatGlmer` functions return output objects of the same class as `lmer` and `glmer`, these outputs can be further used in complement analyses implemented in companion packages of *lme4*, for example, *RLRsim* [17] and *lmerTest* [18] R packages for inference procedures.

We have implemented three features in *lme4qtl* to adapt the mixed model framework of *lme4* for QTL mapping analysis. First, we introduce the positive-definite covariance matrix *G* into the random effect structure, as described in [19, 11]. Provided that random effects in *lme4* are specified solely by *Z* matrices, we represent *G* by its Cholesky decomposition *LL^T^* and applied a substitution *Z* = ZL,* which takes the *G* matrix off from the variance of the vector *u*

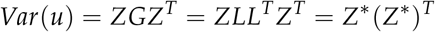

Second, we address situations when *G* is positive semi-definite, which happen, for example, if genetic studies include twin pairs [1]. To define the *Z\** substitution in this case, we use the eigendecomposition of G. Although *G* is not of full rank, we take advantage of *lme4’* special representation of covariance matrix in linear mixed model, which is robust to rank deficiency [16, p. 24-25].

Third, we extend the *lme4* interface with an option to specify restrictions on model parameters. Such functionality is necessary in advanced models, for example, for a trait measured in multiple environments (Supplementary Note 1).

We note that the later two features are available only in *lme4qtl*, but not in other *lme4*-based extensions such as the *pedigreemm* package [11].

### 2.3 Analysis of the GAIT2 data

The sample from the Genetic Analysis of Idiopathic Thrombophilia 2 (GAIT2) project consisted of 935 individuals from 35 extended families, recruited through a proband with idiopathic thrombophilia [14]. We conducted a genome-wide screening of activated partial thromboplastin time (APTT), which is a clinical test used to screen for coagulation-factor deficiencies [20]. The samples were genotyped with a combination of two chips, that resulted in 395,556 single-nucleotide polymorphisms (SNPs) after merging the data. We performed the same quality control pre-processing steps as in the original study [14]: phenotypic values were log-transformed; two fixed effects, age and gender, and two random effects, genetic additive and shared house-hold, were included in the model; individuals with missing phenotype values were removed and all genotypes with a minimum allele frequency below 1% were filtered out, leaving 263,764 genotyped SNPs in 903 individuals available for GWAS. We compared the performances between our package and SOLAR [2, 21], one of the standard tool in family-based QTL mapping analysis.

## 3 Results

We considered three models for the analysis of APTT in the GAIT2 data, namely polygenic, association and gene-environment interaction.

Before conducting the analysis, we organized trait, age, gender, individual identifier id, house-hold identifier hhid variables and SNPs as a table dat. The additive genetic relatedness matrix was estimated using the pedigree information and stored in a matrix `mat`. A polygenic model m1 was fitted to the data by the `relmatLmer` function as follows.

~~~
m1 <- relmatLmer(aptt ^~^ age + gender + (1|id) + (1|hhid), dat,
 relmat = list(id = mat))
~~~

The proportion of variance explained by the genetic effect (heritability) was 0.56, and its 95% confidence interval, estimated by profiling the deviance [16], was [0.45; 0.84].

We further tested whether the genetic effect was statistically significant by simulations of the restricted likelihood ratio statistic, as implemented in the `exactRLRT` function of the *RLRsim* R package [17]. The p-value of the test was below 2.2 × 10^−16^.

For a single SNP named rs1, the update function created an association model m2 from m1 and the anova function then performed the likelihood ratio test.

~~~
m2 <- update(m1, . ^~^ . + rs1)
anova(m1, m2)
~~~

To automate the GWAS analysis, we created an example `assocLmer` function with several options such as different tests of association and parallel computation. By using the `assocLmer` function, we have replicated some loci previously known for APTT [20] (Supplementary Figure 1) applying the likelihood ratio test and running the analysis in parallel on a desktop computer (2.8GHz quad-core Intel Core i5 processor, 8GB RAM).

The GWAS computation time of the association model with two random effects by *lme4qtl* was 7.6 hours. We performed the same analyses, using SOLAR, and observed a computation time 3 fold larger (25.1 hours, Supplementary Table 3). In additional experiments varying the number of fixed and random effects, the *lme4qtl* package was also several times faster than SOLAR (Supplementary Table 3, Supplementary Figure 2), owing to the efficient lme4 implementation of sparse matrix methods. Note that when a model has a single random effect, SOLAR had a option to apply the eigendecomposition trick and substantially speed up the computation (3.8 hours), while this option has not been implemented in *lme4qtl* (6.6 hours).

We then considered an advanced model of gene-environment interaction, an extension of the polygenic model m1, where the variance of genetic random effect is a function of environment (gender). Supplementary Notes 1 and 2 contain details on model specification and numerical results obtained on the GAIT2 data.

We also evaluated how the *lme4qtl* computation time depends on the sparse structure of covariance matrices (Supplementary Figure 3). When artificially reducing sparsity of the genetic relatedness matrix in the GAIT2 data, we found that the time required to fit the polygenic model increases substantially: it becomes an order of magnitude greater once the sparsity changes from the GAIT2 level 0.98 to 0.60.

## 4 Discussion and Conclusions

We have extended the *lme4* R package, a well-established tool for linear mixed models, for application to QTL mapping. The new *lme4qlt* R package has adopted the *lme4*’s powerful features and contributes with two key building blocks in QTL mapping analysis, custom covariance matrices and restrictions on model parameters. To our knowledge, the *lme4qlt* R package is the most comprehensive extension of *lme4* to date for QTL mapping analysis.

Our package has also limitations. In particular, introducing covariance matrices in random effects implies that some of the statistical procedures implemented in *lme4* might not be applicable anymore. For instance, bootstrapping in the update function from *lme4* cannot be directly used for *lme4qlt* models. Furthermore, the residual errors in *lme4* models are only allowed to be independent and identically distributed, and *ad hoc* solutions need to be applied in more general cases, as we showed for the gene-environment interaction model. However, this restriction on the form of residual errors may be relaxed in the future *lme4* releases, according to its development plan on the official website [22]. Also, *lme4qlt* cannot compete with tools optimized for particular GWAS models with a single genetic random effect: *lme4qlt* allows for association models with multiple random effects. Last, *lme4qlt* is mostly applicable to datasets with sparse covariances, while its use in population-based association studies with dense covariances may lead to a considerable overhead in computation time.

In conclusion, the *lme4qlt* R package enables QTL mapping models with a versatile structure of random effects and efficient computation for sparse covariances.

## 5 Declarations

### 5.1 Ethics approval and consent to participate

The GAIT2 study was reviewed and approved by the Institutional Review Board of the Hospital de la Santa Creu i Sant Pau, Barcelona, Spain.

### 5.2 Consent for publication

Not applicable.

### 5.3 Availability of data and material

Source code of *lme4qtl* is available at https://github.com/variani/lme4qtl.

### 5.6 Author’s contributions

A.Z., M.V.S. and H.B. conceived the study; A.Z. implemented the software; A.Z., M.V.S., H.B. and A.M.P analyzed the data; H.A. and J.M.S directed the study; A.Z. and H.A. drafted the manuscript; all authors read and approved the final manuscript.

## 5.7 Acknowledgments

A.Z. thanks Donald Halstead for reading and providing feedback on early drafts of the manuscript.

## 6 Abbreviations

QTL: quantitative trait locus; SNP: single nucleotide polymorphism; LMM: linear mixed model; GWAS: genome-wide association study; APTT: activated partial thromboplastin time; GAIT2: Genetic Analysis of Idiopathic Thrombophilia 2.

